# Synergy of AMPA and NMDA receptor currents in dopaminergic neurons: a modeling study

**DOI:** 10.1101/024653

**Authors:** Denis Zakharov, Christopher Lapish, Boris Gutkin, Alexey Kuznetsov

## Abstract

Dopaminergic (DA) neurons display two modes of firing: low-frequency tonic and high-frequency bursts. The high frequency firing within the bursts is attributed to NMDA, but not AMPA receptor activation. In our models of the DA neuron, both biophysical and abstract, the NMDA receptor current can significantly increase their firing frequency, whereas the AMPA receptor current is not able to evoke high-frequency activity and usually suppresses firing. However, both currents are produced by glutamate receptors and, consequently, are often co-activated. Here we consider combined influence of AMPA and NMDA synaptic input in the models of the DA neuron. Different types of neuronal activity (resting state, low frequency, or high frequency firing) are observed depending on the conductance of the AMPAR and NMDAR currents. In two models, biophysical and reduced, we show that the firing frequency increases more effectively if both receptors are co-activated for certain parameter values. In particular, in the more quantitative biophysical model, the maximal frequency is 40% greater than that with NMDAR alone. The dynamical mechanism of such frequency growth is explained in the framework of phase space evolution using the reduced model. In short, both the AMPAR and NMDAR currents flatten the voltage nullcline, providing the frequency increase, whereas only NMDA prevents complete unfolding of the nullcline, providing robust firing. Thus, we confirm a major role of the NMDAR in generating high-frequency firing and conclude that AMPAR activation further significantly increases the frequency.

## Introduction

Midbrain dopamine neurons predominantly fire in a low frequency, metronomic manner (i.e. tonic) and display occasional high frequency, burst-like episodes [1, 2, 3]. Tonic spiking patterns of the DA neuron are observed in vivo and in isolated preparations (i.e. slices), however they are less regular in vivo [2, 3]. The transition from tonic firing to a burst is stimulus-driven and evokes transient increases in DA release throughout the brain. The computational function facilitated by transitions from tonic firing to a burst in DA neurons has been studied extensively and the preponderance of evidence suggests that this signal provides a value judgment regarding the salience of environmental stimuli [2, 4].

In vivo, bursts in DA neuron firing are not repetitive, but are typically singular events superimposed on the tonic firing pattern [1, 2, 3]. A burst has been classically defined as any group of three or more spikes starting with an interspike interval shorter than 80 ms and ending with an interspike interval longer than 160 ms [1]. Both in vivo and in slices, bursts can be elicited via transient activation of synaptic inputs, and, in particular, N-methyl-D-aspartate (NMDA) receptor activation [5, 6, 7, 8, 9, 10, 11]. Additionally, the blockade of calcium-dependent potassium (SK-type) channels has also been shown to evoke bursting [12]. However, in contrast to in vivo observations, the SK blockade results in repetitive bursting, where high-frequency episodes are separated by pauses. The repetitive bursting phenomena are also observed following bath application [8, 9], but not iontophoresis of NMDA [10, 11]. Thus, high-frequency spiking can be clustered into repetitive bursts or, more commonly, a single and transient burst. The mechanisms in repetitive bursting that create pauses between bursts have been studied separately from high frequency spiking itself [8, 13, 14]. However, high-frequency spiking is likely to increase DA efflux above background levels and thus capable of providing a signal that alters the computational properties of afferent networks. Therefore it is important to focus on the mechanism of high-frequency firing more generally, which is done in the current study.

SK channels are partially responsible for repolarizing the cell after a spike, as they evoke a long-lasting hyperpolarization after each spike, which extends the interspike interval [15]. Likewise, blockade of this current increases the firing frequency by shortening the interspike interval [12]. This type of high-frequency firing, however, rapidly transitions to depolarization block within the bursts [9, 12]. The mechanism whereby episodes of highfrequency firing are evoked by NMDAR activation are diffcult to explain because SK current activation should still repolarize the neuron leading to long interspike intervals. One possibility is that NMDAR current counteracts the SK current, but then any depolarizing current should evoke high frequency firing, which is not true. For example, a tonic applied somatic depolarization should elevate the firing frequency to the levels observed during bursting in vivo, but it does not [16]. In particular, applied depolarization can increase firing frequency only up to 10 Hz, whereas it reaches above 20 Hz in the bursts.

Stimulation of -amino-3-hydroxyl-5-methyl-4-isoxazolepropionate (AMPA) receptors, which are activated by glutamate like NMDA, elicits quite a different response in the DA neuron. A number of experimental studies suggest that stimulation of NMDA receptors evokes a burst of high-frequency firing, whereas AMPA receptor activation evokes modest increases in firing [5, 6, 10, 11, 17] (but see [18, 19]). The degree of the frequency increase is important because it separates the phasic and background release of dopamine, and the gap between the frequency ranges (below 10 and above 20 Hz) allows for their clear distinction.

To explore how NMDAR activation is capable of evoking higher firing frequencies than observed following AMPAR activation or applied depolarization, we studied the interaction between the currents and the oscillatory mechanism of the DA neuron. Based on the identification of ion channels [12, 20, 21, 22, 23, 24, 25, 26, 27, 28, 29, 30] and modeling studies [13, 31, 32], the maintenance of tonic firing is shown to rely on the interactions of voltage gated *Ca*^2+^ and calcium-dependent *K*^+^ currents, which periodically bring the neuron to the spike threshold and generate metronomic firing activity. In contrast, spike-producing currents (fast sodium and the delayed rectifier potassium) play a mostly subordinate role, adding a spike on top of the oscillations without significant changes to the period or shape of voltage and calcium oscillations [32]. This mechanism is called a subthreshold *Ca*^2+^ − *K*^+^ oscillatory mechanism. In our previous studies [33, 34, 35], we have further shown that the same oscillatory mechanism can produce high-frequency oscillations of the membrane potential in interaction with the NMDAR current. The activation of the NMDAR current increases the frequency of the oscillation by limiting the amplitude of *Ca*^2+^ concentration: *Ca*^2+^ removal is a slow process, and the smaller the change required, the shorter is the interspike interval. The NMDAR current counteracts the SK current without blocking the *Ca*^2+^ − *K*^+^ oscillations due to its magnesium block at low voltages. In contrast, the AMPAR current is strongest at low voltages and, together with opposing the SK current, blocks oscillations all together.

While the effects of the NMDA and AMPA receptor on DA neuron firing are clearly different, they are both activated by glutamate. Considering this, is it difficult to reconcile the apparently opposing effects that AMPA and NMDA have on DA neuron firing when co-activated. The purpose of the current study is to determine the combined effects of AMPA and NMDA on DA neuron firing and, specifically, how co-activation leads to a transition from tonic firing as typically defined (i.e. 1-4 Hz) to high frequency firing (> 20*Hz*).

There are several conductance-based models of the DA neuron [13, 31, 36, 37, 38, 39], including our own [33, 34, 40, 35]. The earlier studies focused on the oscillation underlying the on-off pattern of repetitive bursting either during NMDA [13] or an SK current blocker [31]. The study closest to ours also considers co-activation of the AMPA and NMDA receptors. In this study, the same bursty input acts on both receptors, which leads to their pulsatile activation, especially for the short-lived AMPAR. In contrast, we focus on the situation in which the receptors are activated tonically either by nonbursty asynchronous activity of the glutamatergic inputs in vivo, or by long-lasting application of glutamate in vitro. In this situation, the model by Canavier and Landry [37] would not differentiate the responses to AMPA and NMDA receptor activation: the frequency increase would be similar. Thus, the question whether AMPAR co-activation further increases the frequency or obscures firing by quickly inducing depolarization block did not emerge.

In the current study, we initially chose a minimal model to avoid the complexities of conductance-based models, which has been used previously to study DA neuron dynamics [41, 42]. While conductance based models are critical to enumerate the potential factors contributing to the major biophysical properties of the DA neuron, minimalistic models are necessary to find the most essential ones. The major biophysical properties of the DA neuron are:

- Tonic low-frequency (1-4 Hz) oscillations in isolation;
- Elevation in the frequency during NMDA receptor stimulation above 20 Hz;
- Frequency during AMPA receptor stimulation or depolarizing current injection below 10 Hz.

The model is based on the classical FitzHugh-Nagumo oscillator [43] and includes the nonlinearity of the SK-type calcium-dependent potassium current, which allows for the above properties to be reproduced. Thus, the subthreshold calcium-potassium mechanism for oscillations, implemented even in the abstract form of FHN model, is responsible for the major properties of the DA neuron. We analyze dynamical mechanisms in the minimal model and further simulate the same results in our biophysical model [35] for better quantitative predictions.

The models differentially respond to AMPA and NMDA tonic synaptic inputs, however, we now consider their co-activation. We show that, combined with NMDA, AMPA may further increase the frequency of firing. The peak frequency that can be achieved with the co-activation of the receptors is 20% higher than for the activation of the NMDA receptor alone in the minimal model, and reaches 40% in the biophysical model. Thus, our results confirm the hypothesized major role of NMDA receptors in generating high frequency firing, as observed in experiments. Further, we show that AMPA receptor activation may contribute to bursting (increasing the frequency in the burst), or obscure the burst by inducing depolarization block.

## 1 Models

### 1.1 Minimal model

First, we study dynamical mechanisms in the following minimal neuron model based on the classical FHN oscillator [43]:

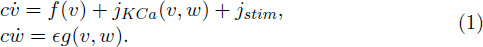

As in FHN oscillator, the first variable *v* describes membrane potential, and functions *f*(*v*) is set as

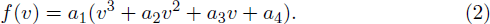

The second variable w in (1) represents a slow recovery process, such as gating of a repolarizing current. The nonlinear term *j*_*KCa*_(*v, w*) represents such a current and provides feedback. This term replaces a linear function of *w* in the classical FHN oscillator. As we have shown before [41, 42], this substitution provides the major biophysical properties of the DA neuron in the model. The nonlinear function represents an SK-type *Ca*^2+^-dependent potassium current [13]:

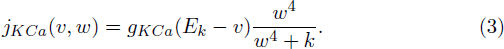

Here, all parameters are made dimensionless to match the formalism of FHN model (see below). *g*_*KCa*_ is proportional to the maximal current density, *E*_*K*_ is the *K*^+^ reversal potential, *k* is the half activation of the current. Accordingly, variable *w* corresponds to calcium ion concentration, and *ϵ* is a small parameter that makes the variable slow and reffects calcium bufiering (see below).

Above the *v* axis in the phase plane, the *v*-nullcline is *N*-shaped (see e.g. Fig. 4). But due to the nonlinearity introduced by the SK-type current (3), it has another folded branch below the axis: *v*-nullcline is symmetric about the *v* axis (not shown). Intersections with the *w*-nullcline in that region may become stable equilibrium states and attract trajectories even from the region of positive *w*. To avoid that, we choose function *g*(*v, w*) (and, thus, the *w*-nullcline *v* = *g*(*v, w*)) to be piece-wise linear

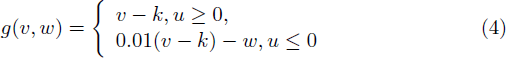

We further introduce the parameter *c* (*c* = 1.1 *\** 10^−4^) in the model to calibrate the time and scale the frequency to be in Hz. The rest of the variables are dimensionless and are calibrated to be of the same order of magnitude and within a unit interval, which is optimal for mathematical analysis and simulations. The recovery variable can be interpreted as calcium concentration since we used a nonlinearity of the calcium-dependent potassium current (3). Thus, the function *g*(*v, w*) takes into account calcium currents. Given the abstract cubic function *f*(*v*) from the classical FHN system, the model does not include particular currents (e.g subthreshold or spike-producing). The cubic function *f*(*v*) is independent of *g*(*v, w*), meaning that not only the calcium currents, but also other depolarizing currents are taken into account.

The minimal model (1) includes two external excitatory stimuli: AMPAR and NMDAR synaptic currents. They are represented by the corresponding terms in *j*_*stim*_:

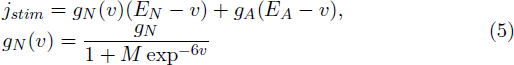

where *g*_*A*_ and *g*_*N*_ are dimensionless maximal conductances of AMPAR and NMDAR channels respectively, *E*_*N*_ and *E*_*A*_ are their reversal potentials. The AMPAR conductance is independent of the voltage, and the conductance of the NMDAR is *g*_*N*_ (*v*), which was modified from [13]. The dimensionless parameter M captures the dependence of NMDAR on the concentration of magnesium, which blocks the receptor at low voltages and, thus, creates the nonlinearity.

We fixed all parameters (*a*_1_ = *−*1, *a*_2_ = 1.35, *a*_3_ = 0.54, *a*_4_ = 0.0539, *k* = −0.585, *M* = 0.2, *E*_*N*_ = *E*_*A*_ = 0, *g*_*KCa*_ = 0.5, *E*_*KCa*_ = −1, *k* = 10, ε = 0.01) except for *g*_*A*_, *g*_*N*_, which vary as the control parameters of the system (1). For this parameter set, the model (1) without an external stimulus demonstrates robust low-frequency tonic (periodic) oscillations.

### 1.2 Biophysical model

Second, we simulate the same results in our biophysical model [35] for more quantitative predictions. Briefly, the model consists of three equations for the voltage, intracellular *Ca*^2+^ concentration and the gating variable of the ERG respectively:

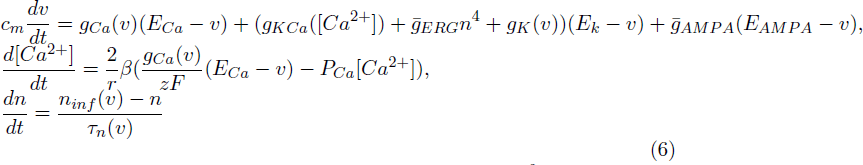

In the voltage equation, *g*_*Ca*_(*v*) is a voltage-dependent *Ca*^2+^ conductance; *g*_*KCa*_(*v*) is a *Ca*^2+^-dependent *K*^+^ conductance; *g*_*ERG*_ is the ERG conductance. A small leak conductance *g*_*l*_ and a voltage-gated instantaneous potassium conductance *g*_*K*_ are included to limit the input resistance and the voltage, respectively. The spike-producing currents were found not to be essential for generating either high- or low-frequency activity [11, 32]. Thus, we calibrated the model to reproduce the voltage oscillations when the fast sodium current is blocked. The presence of the ERG current is one of the factors that make the biophysical model much more complex than the minimal one. The equation for the gating variable of the ERG current in system (6) is in the standard Hodgkin-Huxley form with the following activation function and time constant:

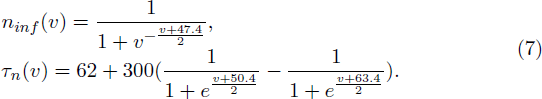

The parameters are a product of calibration of this current to sustain pacemaking in the absence of the *Ca*^2+^-dependent potassium current. The voltage dependence of the *Ca*^2+^ current

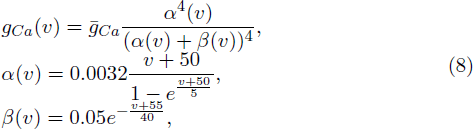

replicates the characteristic low-threshold L-type current found in DA neurons.

The fourth power dependence of the SK type *Ca*^2+^-dependent potassium conductance on *Ca*^2+^ concentration

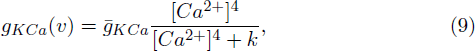

is typically used to best represent the characteristics of the current. This is exactly the same dependence (3) as in the minimal model, but in dimensional variables.

The additional potassium current has the conductance 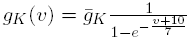.

As in the minimal model, the conductance 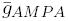 is a constant density of the AMPAR current, but now it is in *mS/cm*^2^ as well as all other conductances in the biophysical model. Also very similar to the minimal model, the nonlinear function of the voltage

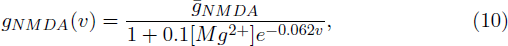

reflects the activation of the NMDAR current. Here, [*M g*^2+^] is magnesium concentration, and the magnesium block is treated as instantaneous.

The calcium equation in system (6) corresponds to the *w* equation in the minimal model (1). It represents balance between *Ca*^2+^ entry via the L current and *Ca*^2+^ removal via a pump. After entering the cell, *Ca*^2+^ binds to a buffering protein. Buffering is assumed instantaneous and is taken into account by the buffering coefficient *β*. This coefficient is the ratio of free to total intracellular *Ca*^2+^. Therefore, [*Ca*^2+^] represents free intracellular *Ca*^2+^ concentration.

All parameter values were the same as in our previous publication [35], except for the conductance of the ERG current, which was lowered to *g*_*ERG*_ = 2. See [35] for more details on the model construction and calibration.

### 1.3 Temporal profile of synaptic receptor activation

In both models, synaptic receptors are activated in a tonic fashion with no kinetics. The reason is that we reproduce experimental results obtained in vitro, in which the timescale of receptor stimulation is much longer than their activation timescale. For example, a typical iontophoresis pulse lasts for 200 msec (e.g. [11]), whereas AMPA and NMDA receptors activate within 1 and 7 msec respectively. The tonic or long-pulse activation of the receptors in vitro mimics tonically active synaptic inputs or long pulses of their activation in vivo. Thus, along with in vitro experiments, our modeling is also relevant to tonic background activation as well as modulation of synaptic inputs in vivo at the timescale of tens or hundreds of milliseconds.

### 1.4 Firing frequencies

In accord with the electrophysiological literature on DA neurons, the models build on the well-established calcium-potassium mechanism for oscillations [1, 32, 15] and mimics other (e.g. spike-producing) currents that are subordinate to the oscillatory mechanism [21, 22, 32, 44]. The biophysical model has been tested and validated in our previous publications [33, 34, 35]. The papers show that the spike-producing currents can be omitted in the model since they only slightly change the frequency.

This approach is similar to that in the integrate-and-fire model [45, 46], where a spike is registered whenever the voltage reaches a threshold. However, the blockade of the spike-producing currents in DA neurons does not change the period and the shape of oscillations between the spikes much [32]. To reflect that in the model, the voltage should reset not to the minimum, but back to the threshold so that the oscillation continues unperturbed. Thus, the firing frequency is measured as the frequency of intersecting the spike threshold in the model (−0.4 in the minimal model, and −40*mV* in the biophysical one). We connect our results with experiments by quantitative comparison to the firing frequencies measured in different conditions.

## 2 Results

### 2.1 Responses of the model to co-activation of synaptic currents

The case of separate influence of the AMPAR and NMDAR synaptic currents was studied in [41, 42]. The AMPAR current increases firing rate of the DA neuron but at the same time quickly and monotonically decreases the amplitude of voltage oscillations resulting in their localization in the subthreshold range and cessation of firing for small increase in the AMPAR conductance (Fig. 1A). The NMDAR current does not decrease the amplitude of voltage oscillations and mostly influences the frequency (Fig. 1B). As NMDAR conductance grows, the frequency increases first, attains a maximum at an intermediate value of the conductance and then gradually decreases (Fig. 2A NMDA ONLY). Since the DA neuron may transition to repetitive bursting at strong NMDAR activation, and this mechanism is not in the model, we do not consider the further increase of NMDAR current. However, the biphasic frequency modulation by NMDA has been reported in an experimental study [48].

**Figure 1:**
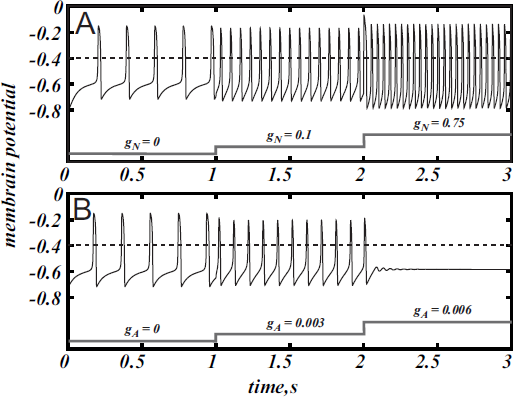
NMDAR current increases the firing frequency above 20 Hz (A) whereas AMPA evokes a much smaller frequency increase and provokes depolarization block (B) in the minimal model of DA neuron (1). Spikes a registered as the voltage oscillation crosses the threshold of -0.4 (dimensionless units).

**Figure 2:**
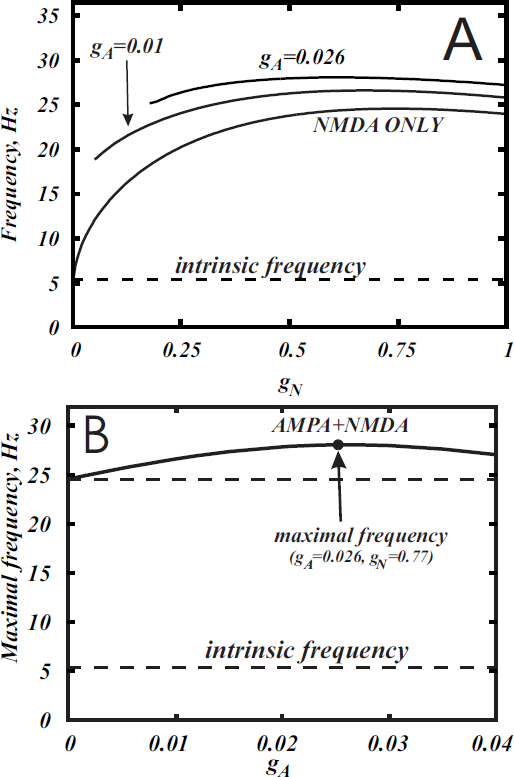
Co-activation of AMPA and NMDA receptor courrents allow one to reach a significantly higher frequencies. (A) Dependence of the frequency in the minimal model (1) on the conductance of NMDA current *g*_*N*_ for the different values of *g*_*A*_. The curves correspond to horizontal slices of the Fig 4. (B) The maximum frequency calculated over the whole range of *g*_*N*_ for each value of *g*_*A*_. The additional frequency growth exceeds 20%. The intrinsic frequency in the graph corresponds to the tonic firing without any external stimulus.

Here we consider in detail combined influence of AMPA and NMDA receptor currents. As well as without NMDA, for the small values of *g*_*N*_ and large enough values of *g*_*A*_, the neuron is not firing (Fig. 1B). Interestingly, increasing NMDAR current rescues firing, and this results in firing frequencies greater than those for NMDAR activation only. Thus, it is possible to obtain a higher frequency in the case of combined influence of both synaptic currents than in the case with the NMDAR current only (Fig. 2A, 5B, 7A). We obtain such elevated frequency in a wide range of *g*_*A*_ and *g*_*N*_. The highest frequency is achieved for *g*_*A*_ = 0.026, *g*_*N*_ = 0.77 (Fig. 2B). In this minimal model, co-activation of the receptors adds up to 20% to the frequency.

The frequency distribution diagram in Fig. 3 shows a large region of high frequency activity (orange and yellow). The frequency is calculated at the intersection of the voltage oscillation with the spike threshold. Thus, the region where the voltage oscillations reduce in amplitude and remain in the subthreshold range are truncated (e.g. near Andronov-Hopf bifurcation; see, for example, Fig.4B). As AMPAR conductance is increased, the firing region shrinks and a greater part of the frequency dependence on the NMDAR conductance is cut off (Fig. Fig. 2A). NMDAR activation is required to restore firing, and the higher the AMPAR conductance, the greater NM-DAR current is needed. On the other hand, the frequency dependence on NMDAR conductance does not shift much: its maximum stays at a virtually the same conductance. Thus, increasing AMPAR conductance cuts the rising part of the frequency dependence first, and then the region of high firing frequencies all together (Fig. 3). We predict that, depending on the background AMPAR activation in experiments, an abrupt transition from silence to high-frequency firing may be observed instead of a gradual frequency increase as NMDAR is activated. At excessive AMPAR activation, the high frequencies may be abolished all together.

**Figure 3:**
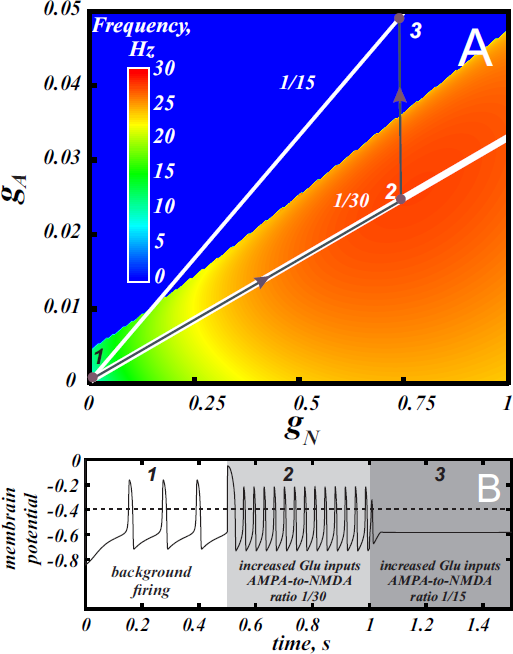
Different combinations of AMPA and NMDA receptor currents may give high frequency firing or depolarization block. (A): Frequency distribution on the parameter plane of AMPA and NMDA receptor conductances. Red region correspond to the high frequency firing. The white lines indicate the conductance ratios 1/30 and 1/15 that correspond to control and facilitation of AMPA current following application of a drug with high abuse potential (see Discussion). The gray arrows and voltage traces in panel (B) illustrate increase in the firing frequency by greater activation of NMDA and AMPA receptors (2) and then blockade of firing by plastic increase in the AMPA conductance (3) at a longer timescale following drug use.

**Figure 4:**
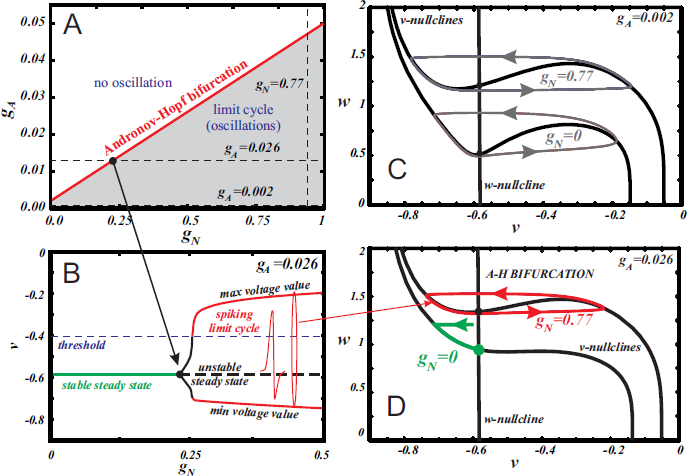
Bifurcation analysis explains the influence of the different combinations of AMPA and NMDA receptor currents. The state diagram (A) and one-parameter bifurcation diagram (B) of the minimal model (1). Panel (B) illustrates the transition along the horizontal dashed line at *g*_*A*_ = 0.026 in panel (A). Panels (C) and (D) show the evolution of the *v*-nullcline in the model phase plane. At a low AMPAR activation (C), a limit cycle of oscillations is initially present. With growing *g*_*N*_, the *v* nullcline flattens, but does not unfold, which preserves the oscillations and increase their frequency. At a high AMPAR activation (D), the equilibrium state is initially stable. With growing *g*_*NMDA*_ the minimum of the *v*-nullcline moves to the left with respect to the *w*-nullcline. As a result, the stable steady state loses its stability and a limit cycle emerges through a supercritical Andronov-Hopf bifurcation. The frequency of these oscillations is highest because the amplitude with respect to the slow variable *w* is minimal.

### 2.2 Dynamical mechanism of the high frequency activity

Next, we use our minimal model to explain the dynamical mechanism of the frequency increase and cessation of firing. The model has the minimal number of variables required to reproduce oscillations: its phase space is a plane of v and w. Thus, we can use nullcline analysis and give a graphic explanation of the synergy between the NMDAR and AMPAR synaptic inputs. In [41, 42] we discussed separate influence of both synaptic currents and found simple geometric explanation to the distinct responses they elicit. The NMDAR current increases the frequency due to flattening (but not unfolding, Fig. 4C,D) of the *v*-nullcline. This change in the geometry of the nullcline determines a decrease in the amplitude of the slow variable *w*, and, consequently, directly reduces the period. By contrast, the AMPAR current mostly shifts the folded region of the *v*-nullcline to the right, and, consequently, stabilizes the equilibrium state via Andronov-Hopf bifurcation. Thus oscillations are lost already at small values of the conductance of the AMPAR current.

Here we further observe that oscillations are preserved for a higher AMPAR conductance if NMDAR is active together with AMPAR (see Fig. 3). The equilibrium state which was stable at high AMPAR and low NMDAR conductances becomes unstable with increasing *g*_*N*_ via a supercritical Andronov-Hopf bifurcation, and a stable limit cycle emerges (Fig. 4A,B). If the limit cycle intersects the threshold (Fig. 4B), it corresponds to neuron firing. With further elevation of *g*_*N*_, the firing frequency increases. To understand how NMDAR activation rescue firing, we conduct nullcline analysis in the minimal model. At a moderate NMDAR conductance, the distance between the minimum of the voltage nullcline and the calcium nullcline is greater than at no NMDAR activation (Fig. 4C,D compare *g*_*N*_ = 0 and 0.77). This is why the Andronov-Hopf bifurcation on Fig. 3A,B occurs at greater values of the AMPAR conductance as NMDAR conductance grows.

Simultaneously, AMPAR activation contributes to the unfolding of the voltage nullcline (Fig. 5), and increases the frequency further (Fig. 2). Note that the observation that oscillations are more robust in NMDA has been made for our detailed biophysical model in [35]. Further, robustness of highfrequency oscillations in NMDA was also observed in experiments, but with respect not to AMPA, but to GABA receptor activation [49].

**Figure 5:**
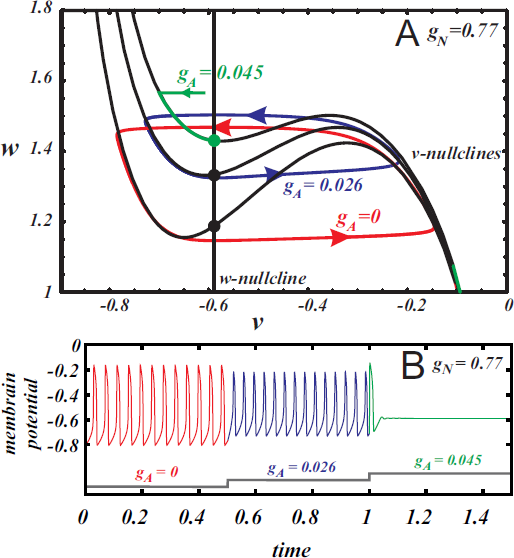
The nullcline analysis explains the synergy of AMPA and NMDA. (A): the evolution of the *v* nullcline and the limit cycle in the phase plane as AMPAR current is introduced. The three cases are no AMPAR activation (red), maximal frequency case (blue) and suppression of oscillations (green). (B): The voltage traces corresponding to the different states represented in the phase plane.

### 2.3 Quantitative prediction of the frequency growth

Here we return to our biophysical conductance-based model (6) to confirm our results and make better quantitative predictions. The frequency was calculated at the intersection of the voltage trace with the spike threshold of -40 mV. Fig. 6 shows the firing frequency as a function of NMDAR conductance for different fixed values of AMPAR conductance. The model displays qualitatively similar behavior as the minimal model. In particular, separately, NMDAR activation increases the firing frequency up to 50Hz, whereas AMPAR does not. Co-activation of AMPAR together with NMDAR leads to a further frequency growth. We found that the additional frequency increase is greater in the biophysical model than in the minimal model and it can be more than 40%. The greater frequency increase is due to differences in the shape of the nullclines better capturing the biophysical characteristics of the currents. The biophysical model is also more accurate at higher NMDAR activation, where a long interval of frequency decrease is truncated by blockade of firing. Again, this is due to the shape of the nullclines. I particular, the *Ca*^2+^ nullcline has sigmoid shape, which forms a stable steady state correspond to depolarization block. With stronger AMPAR activation, the amplitude of voltage oscillations decreases (similar to the minimal model; data not shown) and confine in the subthreshold range. Thus the simulated neuron stops firing. Further, the steady state of depolarization block becomes stable and subthreshold voltage oscillations disappear as well. Because AMPAR current alone leads to depolarization block already at very low values, activation of NMDAR is required to rescue the subthreshold voltage oscillations and firing first, and only then increases the frequency (Fig. 7). This feature is also well captured in the minimal model (Fig. 2A) and these models have only a difference in the velocity of frequency growth with the elevation of the NMDA current: the firing rate in the minimal model grows faster. Thus, the reduced and the conductance-based models demonstrate qualitatively similar firing frequency growth during co-activation of the AMPAR and NMDAR currents (compare voltage traces in Fig. 7).

**Figure 6:**
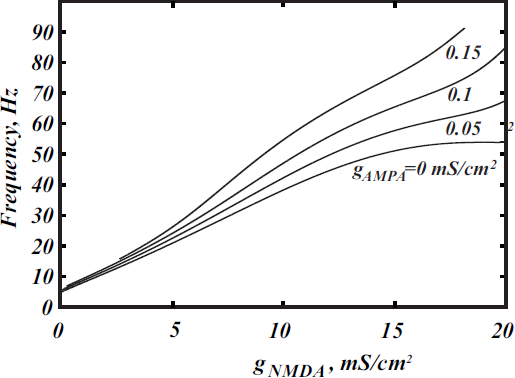
Higher frequencies are achieved for NMDAR and AMPAR current co-activation in the biophysical model (6). The dependence of the frequency on *g*_*NMDA*_ is shown for the different values of *g*_*AMP*__*A*_. The frequency is calculated at the intersection with the threshold -40mV.

**Figure 7:**
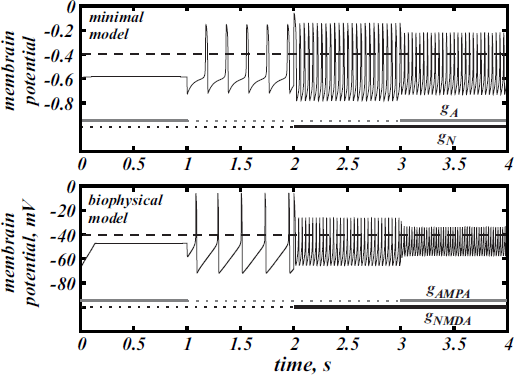
The voltage traces for the minimal (1) and biophysical (6) models under the action of AMPAR and NMDAR currents. In the minimal model *g*_*A*_ = 0.026, *g*_*N*_ = 0.77. In the biophysical model *g*_*AMP*__*A*_ = 0.16, *g*_*NMDA*_ = 8.

## Discussion

In this paper, we consider the co-activation of AMPAR and NMDAR synaptic currents on a DA neuron. Previous data and simulations suggest that these receptors differentially affect DA neuron firing: NMDAR increases the firing frequency while AMPAR causes depolarization block. Therefore, one may expect these effects to oppose one another, where an increase in the frequency caused by NMDAR is impeded by the AMPAR as these receptors are both activated by glutamate and, therefore, should activate together. Thus, the high frequencies evoked by NMDAR activation may not be achievable in vivo, and the effect of receptor co-activation is unclear. Resolving this issue was the focus of the current study.

The simulations presented herein demonstrate that NMDAR activation increases firing frequency with further increases following AMPAR activation. The additional frequency increase mediated by the AMPAR is consistent in both models and exceeds 40% in the biophysical model [35]. However, it is only possible for certain conductances of the AMPAR and NMDAR. In particular, there is an intermediate range of NMDAR activation, in which moderate AMPAR activation further increases the frequency. At either lower or higher NMDAR activation, the AMPAR also increases the firing frequency, but overall increase remains low (*<* 20 Hz, Fig. 6). At any NMDAR conductance, if there is excessive AMPAR activation, this suppresses the *Ca*^2+^ *− K*^+^ mechanism for voltage oscillation all together, sends the neuron into depolarization block, and eliminates firing. However, the stronger the NMDAR activation, the higher the AMPAR conductance is required for depolarization block. Below this value, the growth of AMPAR conductance leads to increases in DA neuron firing. Experimental data support this result as NMDAR and AMPAR co-activation has been shown to increase [50] the firing frequency of the DA neuron.

The effect of co-activation of AMPA and NMDA receptors on DA neurons has been investigated in previous modeling studies where synergistic increases in firing were observed [37]. In addition, [47] show that tonic activation of NMDA receptor delays entry into depolarization block, whereas AMPA does not. Our models shows exactly the same properties and predicts synergistic effect of these receptors on the frequency growth. Note that we consider not transient, but steady frequencies. DA neurons display a spike frequency adaptation, and the transient frequencies are somewhat higher. This is an additional factor that can contribute to burst episodes observed in experiments, and this is not considered in the current paper. Similar to the currently proposed model, Canavier and Landry [37] demonstrated that concurrent activation of AMPA and NMDA receptors increases the firing frequency above that evoked by stimulation of NMDA receptors alone. However, we predict a much greater frequency increase, which could be attributable to the different mechanism proposed here than in [37]. We show how a tonic level of AMPAR activation may increase the firing frequency by flattening the voltage nullcline in the model, whereas in [37] AMPAR activation is short and works by transient opposition to the hyperpolarizing currents. The mechanisms of the frequency increases are not mutually exclusive, however, and could be synergistic in vivo. For instance, asynchronous glutamatergic inputs may summate to yield tonic background activation of AMPA and NMDA receptors and give a moderate increase in the firing frequency. A burst may reflect a further transient frequency increase in the DA neuron due to pulsatile stimulation, which requires transient synchronization of the inputs (by [37]), or simply greater tonic activation of the receptors, which requires increase in firing of the glutamatergic neurons (by our mechanism).

The effect of AMPA depends on the absolute value of the NMDAR conductance, as well as the AMPA-to-NMDA current ratio. Interestingly, many drugs of abuse alter this ratio so that the AMPAR becomes stronger relative to the NMDAR [51]. Taken together with overall excitatory influence of drugs of abuse on DA neurons, one would expect that, in their presence, the greater AMPAR contribution facilitates high-frequency bursts of the DA neuron compared to the control conditions. However, our results show that alterations in the AMPA-to-NMDA ratio may only impede the ability to evoke high-frequency bursts. Our calculations in the biophysical model [35], show that the current ratio measured in experiments (0.4 [51]) in response to a short voltage pulse corresponds to the conductance ratio around 1/30. The solid white line labeled 1/30 in fig. 3 corresponds to this ratio of the AMPA/NMDA conductances. Interestingly, the line passes through the maximum of the frequency distribution. Thus, in the control conditions, the AMPA-to-NMDA ratio is optimal to evoke high-frequency firing. The white line labeled 1/15 in Fig. 3A corresponds to the AMPA-to-NMDA ratio doubled, which mimics the facilitation of AMPAR observed following application of drugs with high abuse potential ([51]). This line is mostly outside of the oscillatory region, meaning that co-activation of AMPA and NMDA receptors will lead to a blockade of firing and possibly an extended state of depression of the DA neuron. The graph shows that the decrease in NMDAR and AMPAR conductances along the line will restore firing, but the maximal frequency may only be a half of that achieved in the control. Thus, the increase in the AMPA-to-NMDA ratio, which results from the acute application of a drug of abuse, moves the DA neuron away from the optimal conditions to evoke high-frequency bursts.

In summary, the predictions that come from our model fit well with the time course of adaptations in the DA system following drug of abuse. The initial short-term increase in firing through co-activation of AMPA and NMDA could facilitate the prediction of highly salient environmental stimuli such as drugs of abuse and cues that predict them [52, 53] (transition from 1 to 2 in fig. 3). Additionally longer time scale, persistent changes in plasticity (i.e. AMPA/NMDA ratio), which are predicted to reduce firing of the DA neuron, may provide a mechanism for hypodopaminergic states observed following drug use [54] (transition from 2 to 3 in fig. 3). The observed depression in DA neuron firing by changing AMPA/NMDA could also provide mechanism of the decreased hedonic effects of drugs of abuse with repeated use as well as the devaluation of all environmental ambient rewards. Experimental studies will be necessary to test these predictions.

## Acknowledgment

D.Z. was partly supported by the Russian Foundation for Basic Research grant 14-02-00916-a. A.K. and C.L. were partly supported by NIAAA grant R01A022821. B.G. was partly supported by ANR-13-NEUC-0003-01, IdEx ANR-11-0001-02 PSL and LabEx ANR-10-LABX-0087 (France). B.G. acknowledges support from the Federal Competitiveness Program of the National Research University Higher School of Economics (Russia).

